# Evaluating Acridones as Novel Therapeutics for Human Babesiosis

**DOI:** 10.64898/2026.01.07.698299

**Authors:** Pratap Vydyam, Elizabeth Zhang, Anasuya C. Pal, Rozalia A. Dodean, Papireddy Kancharla, Jane X. Kelly, Choukri Ben Mamoun

## Abstract

**Background:** Human babesiosis is an emerging tick-borne disease caused by *Babesia* parasites, most notably *Babesia microti* and *Babesia duncani* in North America and *Babesia divergens* in Europe. Infections can be severe, persistent, or relapse despite treatment, and current therapeutic options remain limited, underscoring the urgent need for new and effective treatment options.

**Methods:** Acridone derivatives, originally developed as potent antimalarial agents against multiple life cycle stages of the malaria parasites, were evaluated for their antibabesial activity using continuous *in vitro* culture systems of *B. duncani* and *B. divergens*. Lead candidates were assessed for selectivity against human cell lines to establish preliminary safety profiles, and select compounds were further advanced into preliminary *in vivo* efficacy studies using murine models of *B. duncani* and *B. microti* babesiosis to assess their therapeutic potential.

**Results:** Nine prioritized acridone derivatives demonstrated potent *in vitro* activity against both *B. duncani* and *B. divergens*, accompanied by favorable selectivity indices relative to human cell lines. However, *in vivo* evaluation of representative compounds did not achieve parasite clearance in murine models. Structure-activity relationship (SAR) analyses highlighted key structural features that are critical for maintaining antibabesial potency and offer guidance for further lead optimization.

**Conclusions:** Acridone derivatives show strong *in vitro* antibabesial activity and represent promising lead chemotypes for therapeutic development. To advance these candidates, future studies focused on optimizing their pharmacokinetic properties, improving in vivo efficacy, and evaluating synergistic combination regimens will be essential for progressing toward effective treatments for human babesiosis.

## Introduction

Human babesiosis is an emerging infectious parasitic disease caused by obligate intracellular protozoan parasites belonging to the Babesia genus. Human babesiosis is known to be caused by nine species of Babesia, of which *Babesia divergens* is the most common cause in Europe, and *Babesia microti* and *Babesia duncani* are the most common cause in the United States [1]. *Babesia spp* are spread through tick bites, but in rare cases, can also be transmitted trans placentally, by blood transfusion, or by organ transplantation. Clinical babesiosis is relatively uncommon, with less than 3,000 cases reported annually in the US. However, the number of babesiosis cases have been increasing over the past 15 years, mostly concentrating in the Northeastern US [2]. Among the nine *Babesia* species known to cause human babesiosis worldwide, *B. microti* is responsible for most clinical cases reported to date and is considered to be endemic in the United States [3–5]. Other cases of human babesiosis include *B. divergens* in Europe and *B. duncani* in western US [6–8]. The increase in the geographic distribution of tick vectors, which has been influenced by the environmental changes of the last decades and various anthropogenic factors, is considered the main driver of the recent increase in tick-borne infections. Over 16,000 cases of human babesiosis in the US were reported to CDC between 2011 and 2019. Annual cases more than doubled during this period [2].

Babesiosis infection in humans causes nonspecific flu-like symptoms, but in immunocompromised individuals, individuals with chronic heart, lung, renal, or liver disease, and in individuals >50 years of age, babesiosis can lead to organ failure and death [9, 10]. There is no vaccine available for human Babesiosis. To treat mild and severe cases of Babesia, the CDC currently recommends a 7 to 10-day course of atovaquone in combination with azithromycin (orally in mild cases and intravenously in severe cases) or mono-therapy with clindamycin or quinine [8]. In highly immunocompromised patients, treatment is continued until the patient has two consecutive weeks of negative blood smears. This prolonged course is problematic due to non-trivial adverse side effects associated with treatment, including diarrhea for atovaquone/azithromycin and an increased risk of *Clostridioides difficile* infection in the case of clindamycin [11]. Furthermore, recrudescence after atovaquone or azithromycin treatment have increasingly been reported and have been traced to drug-resistance mutations [12].

In light of these persistent clinical challenges and the growing incidence of human babesiosis, there is a pressing need for new therapeutic agents validated through both *in vitro* and *in vivo* models. Past work with antimalarial compounds have demonstrated that they can share antibabesial effects, highlighting the biological relatedness of *Plasmodium spp* and *Babesia spp* as a fruitful premise for the discovery of pan-antiparasitic compounds [13, 14]. As part of our ongoing efforts to discover and optimize novel, safe, and mechanically distinct inhibitors of *babesia* parasites, we sought to evaluate a new acridone chemotype, previously characterized for potent antimalarial activities against multiple life cycle stages of the malaria parasites [15–17], for its antibabesial potential. These acridone analogs demonstrated robust oral efficacy in multiple rodent models of malaria, with favorable safety, and metabolic profiles. To date, however, neither antibabesial assessments nor comprehensive structure-activity relationship (SAR) studies of these acridones analogs have been reported.

Here, we report the first evidence for the antibabesial effects of acridone compounds against the major etiological agents of human babesiosis, along with preliminary structure-activity relationship (SAR) analysis. We investigated a panel of optimized acridone compounds as potential drug candidates against *Babesia duncani* and *Babesia microti* in our validated mouse models of babesia infection. Nine prioritized acridone derivatives displayed a broad therapeutic window and high *in vitro* efficacy in parasite clearance in our *in vitro* culture conditions. Furthermore, our findings suggest a potential shared molecular pathway by which these compounds exert activity against both *Plasmodium* and *Babesia spp*. Collectively, this data is suggestive of our candidates as a pan-antiparasitic compounds with significant clinical implications.

## Results

### Selection of acridones for *in vitro* and *in vivo* antibabesial activities

A panel of 19 acridone analogs (Table 1) was strategically selected to cover a broad chemical space within the acridone scaffold class, incorporating systematic variations in functional groups, substitution patterns, and lipophilicity. The selected structural diversity was intended to enable the identification of early SAR trends and to define the key molecular features governing antibabesial potency. Importantly, the selected molecules included several analogs that previously demonstrated potent multistage activity against *Plasmodium* spp., favorable oral bioavailability, and robust metabolic and pharmacokinetic profiles [15–17] thus with features that suggested promising translational potential for antibabesial indications. Using this structurally diverse and biologically enriched library, we aimed to evaluate whether key molecular determinants of antimalarial efficacy would be translated to activity against *babesia* parasites and to identify lead chemotypes capable of exerting broad antiparasitic effects.

**Table 1.**
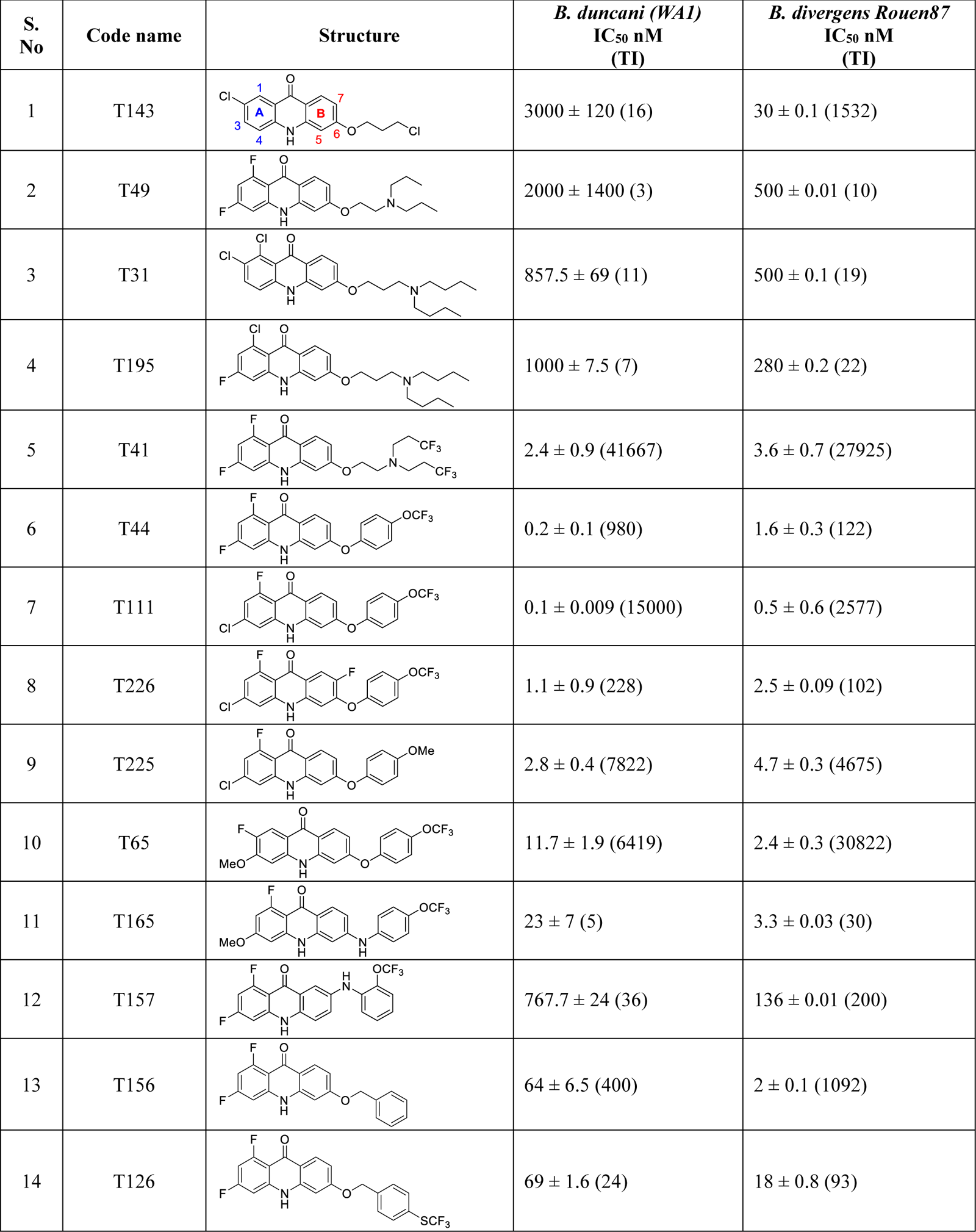
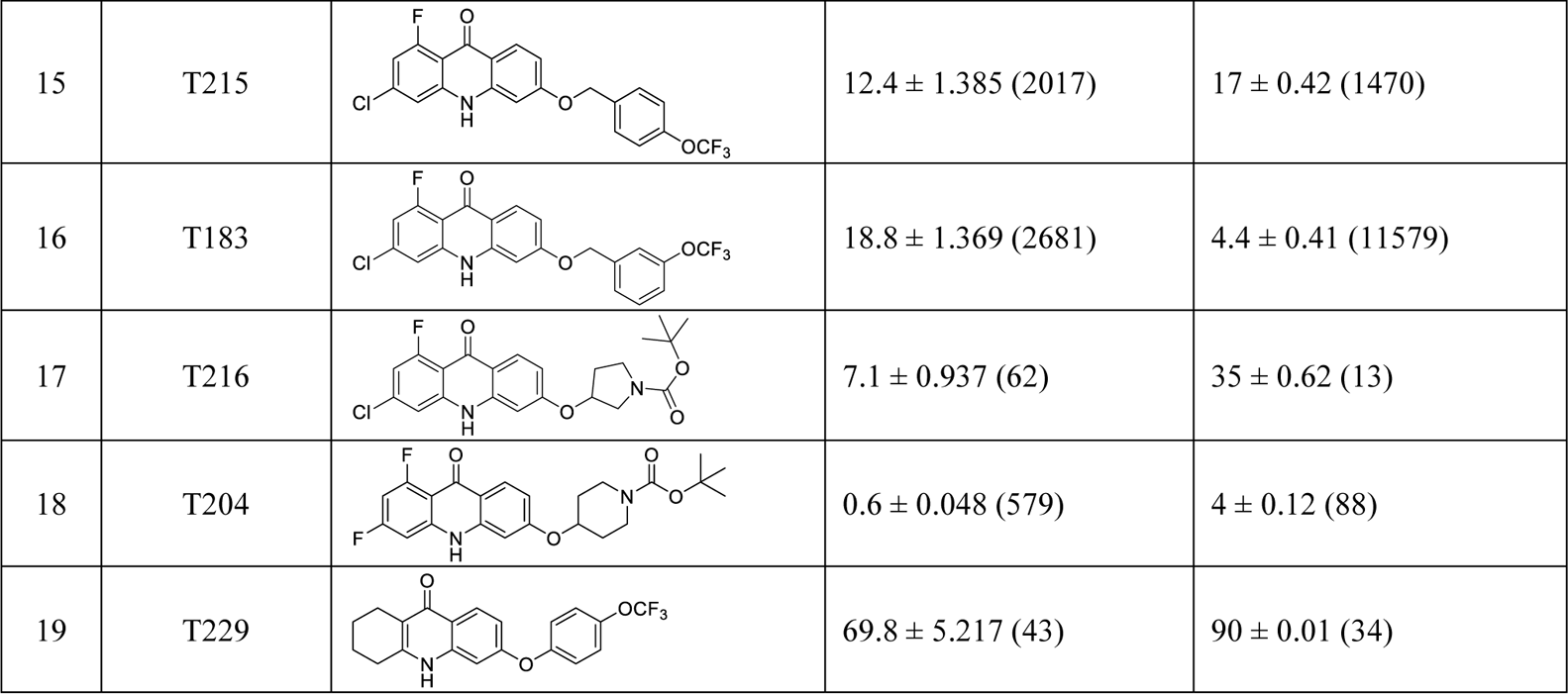
*In vitro* antibabesial activity, selectivity, and SAR of acridone analogs.

### Potency and safety of acridone derivatives against *babesia* parasites *in vitro*

Subsequently, these 19 acridone derivatives were screened for *in vitro* antibabesial activity and cytotoxicity. *In vitro* activity was assessed against continuous cultures of *Babesia divergens* (Rouen 87 strain) and *Babesia duncani* (WA-1 strain) in human erythrocytes, using a SYBR Green-based dose-response assay. Untreated parasites and those treated with 2 µM WR99210 (achieving 100% inhibition) served as controls. At a single concentration of 1 µM, 90% of the compounds exhibited >80% growth inhibition against both *B. duncani* and *B. divergens*, as shown in the heat map (Figure 1A). Dose-response assays were conducted to determine IC_50_ values for individual compounds (Figure 1B and supplemental Figures S1 and S2). Cytotoxicity was evaluated against four pharmacologically relevant human cell lines to calculate therapeutic indices (TI). Of these, nine derivatives T111, T225, T65, T215, T183, T229, T165, T49 and T143, demonstrated potent activity (IC_50_ values ranging from 0.1 nM to 3 µM) (Table 1) and favorable selectivity indices (Table 1). These compounds were selected for further *in vivo* studies due to their high *in vitro* potency against *Babesia* spp., availability, low cytotoxicity, and shared antimalarial activity.

**Figure 1.**
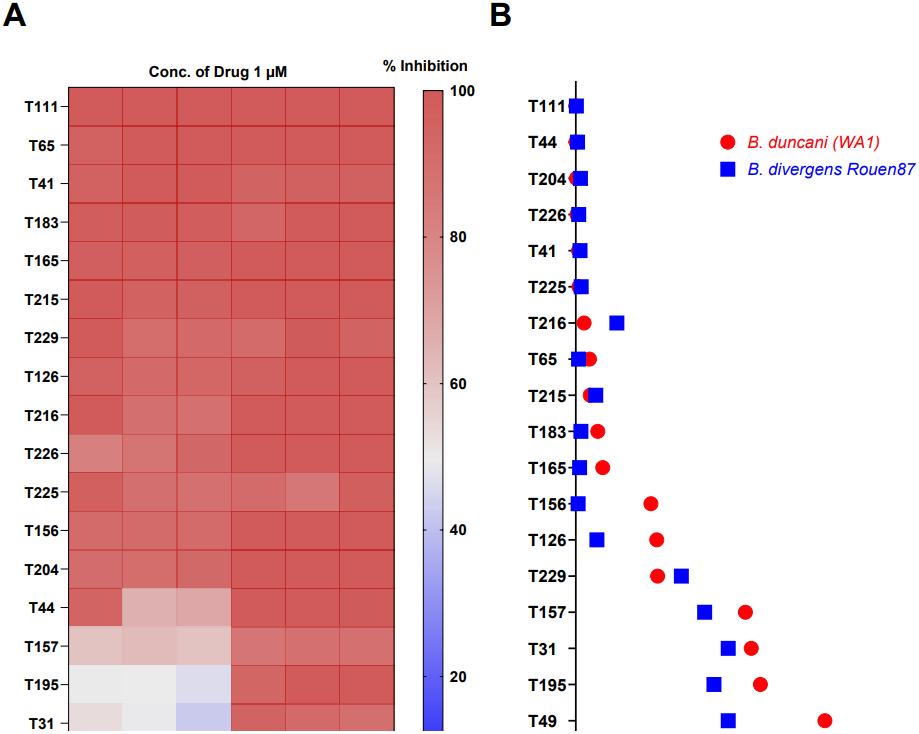
In vitro potency and IC_50_ values of acridone derivatives against *B. duncani* and *B. divergens*. **(A)** Heat map illustrating the *in vitro* potency of 19 acridone derivatives at a fixed concentration (1 µM). The color gradient (red to blue) represents the percentage inhibition of parasite growth for *Babesia duncani* (WA-1 strain) and *Babesia divergens* (Rouen87 strain), with red darker shades indicating higher inhibition. **(B)** Scatter plot depicting the IC_50_ values (in nM) of acridone derivatives against *B. duncani* (red) and *B. divergens* (blue). Compounds are arranged in ascending order of IC_50_ values for *B. duncani*, reflecting their relative efficacy against this parasite.

### Structure-activity relationship (SAR) investigations

*In vitro* activity and SAR investigations of the selected acridones revealed distinct structural features required for achieving optimal antibabesial potency. The first-generation acridones (entries 1–4, Table 1), which contain an alkoxy substituent at the 6-position of ring-B and various halogens on ring-A, showed only moderate activity, with potencies in the low micromolar range against both tested babesia strains. In contrast, incorporation of a terminal trifluoromethyl group on the 6-alkoxy chain, as exemplified by T41 (entry 5, Table 1), resulted in a significant improvement in potency relative to the corresponding first-generation analogs, indicating that terminal trifluoromethyl substitution on the alkoxy chain is highly favorable for antibabesial activity. Notably, most of the second-generation acridones (entries 6–11, Table 1), which feature an aryloxy moiety at the 6-position of ring-B, displayed markedly enhanced *in vitro* potency. These results highlight the pivotal role of 6-aryloxy functionality in driving improved activity. Interestingly, shifting this substitution from the 6-position to the 7-position, as in T157 (entry 12, Table 1), substantially reduced potency, underscoring the positional sensitivity and confirming that substitution at the 6-position is crucial for maintaining antibabesial potency. Replacement of the 6-aryloxy group with benzyloxy moieties (entries 13–16, Table 1) led to a pronounced reduction in potency. However, these analogs remained more active than the first-generation acridones, indicating partial retention of the pharmacophore. Substituting the aryloxy group with cycloalkoxy moieties (entries 17 and 18, Table 1) largely preserved potency, suggesting that certain non-aryl cyclic ether substituents can also be accommodated. Finally, complete hydrogenation of ring-A in the highly potent analog T-111 (entry 7, Table 1) to generate T-229 (entry 19, Table 1) resulted in diminished activity, demonstrating that the aromatic integrity of ring-A is essential for maintaining optimal potency. Overall, the second-generation acridones displayed superior antibabesial activity compared to the first-generation analogs. Nonetheless, additional SAR exploration is warranted to further refine the structural determinants that govern potency within this chemotype.

### In vivo efficacy of acridone derivatives against murine models of human babesiosis

Given their potent *in vitro* antibabesial activity, excellent safety profiles (Table 1), availability, favorable DMPK properties, and previously demonstrated oral efficacy in multiple rodent malaria models [15–17], several promising acridone derivatives, T111, T225, T65, T215, T183, T229, T165, T49 and T143, were selected for *in vivo* efficacy evaluation against Babesia infections in immunocompetent C3H/HeJ mice (n=3 per group, minimum).

Initially, we evaluated the efficacy of T111 against both *B. duncani* and *B. microti* in mouse infection models. Mice were inoculated with 1×10^5^ *B. duncani* or *B. microti* parasites and treated with 40 mg/kg of T111 daily by oral gavage from DPI-3 to DPI-7. In *B. duncani* infected mice, we found no change in the parasite burden or advantage in survival compared to vehicle (PEG400)-treated control mice. (Figure 2A and B). In case of *B. microti* infected mice, the vehicle treated controls naturally cleared the infection between DPI-15 and DPI-20, as expected, and T111 treatment did not significantly reduce parasitemia compared to controls (Figure 2C), indicating no therapeutic benefit in this model. These findings suggest that T111, a potent anti-malarial drug, may require further optimization in terms of dosing and treatment duration to be efficacious against *B. duncani* and *B. microti* infections.

**Figure 2.**
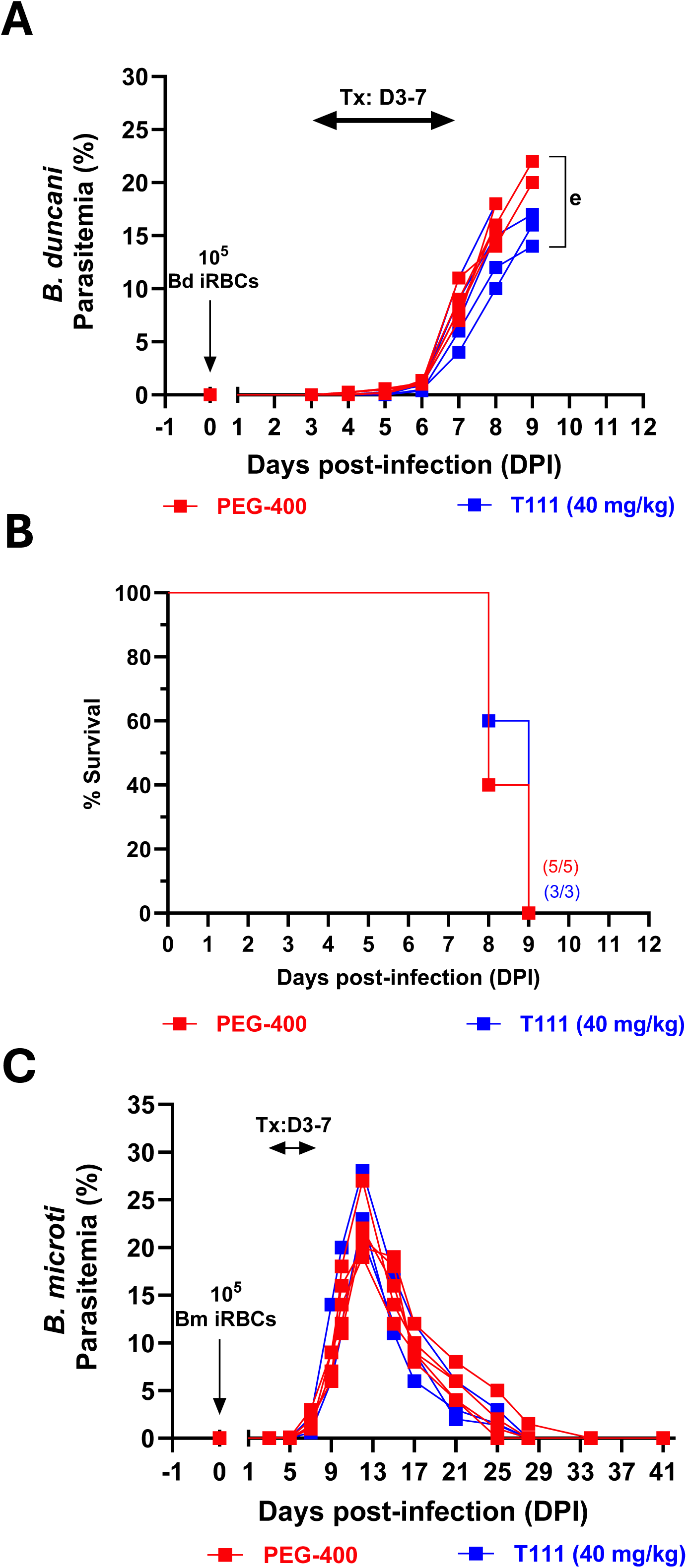
*In vivo* efficacy of the lead acridone derivative (T111) against *B. duncani* and *B. microti* infected mice. **(A)** Parasitemia and **(B)** Kaplan-Meier survival curve of female C3H/HeJ mice (n = 3 per group) infected intravenously with 1 × 10^5^*Babesia duncani* (WA-1 strain)-infected red blood cells (iRBCs), inducing lethal infection. Mice were treated via oral gavage once daily from days post-infection (DPI) 3 to 7 with either PEG-400 (vehicle, red) or T111 (40 mg/kg, blue). e, euthanized. **(C)** Parasitemia in 10^5^ *Babesia microti*-infected red blood cells (iRBCs) inoculated mice treated via oral gavage once daily from DPI 3 to 7 with either PEG-400 (vehicle, red) or T111 (40 mg/kg,). Parasitemia was quantified using Giemsa-stained blood smears, counting a minimum of 3,000 erythrocytes per smear.

Since T111 did not show efficacy against 10^5^ infections dose of *B. duncani*, we further tested efficacy of the remaining shortlisted acridone compounds (T225, T65, T215, T183, T229, T165, T49 and T143) against 10^4^ -*B. duncani*-iRBC infected mice. Depending on the availability of these compounds, infected mice were treated with each compound at 30 mg/kg daily for 5 days, starting on day 1 post-infection (DPI-1 to DPI-5) and parasitemia and survival were monitored until DPI-41. Vehicle (PEG-400) treated controls exhibited peak parasitemia of 10% by DPI-11, with all mice succumbing to infection by DPI-11, consistent with established models (Fig. 3A and K). Atovaquone treatment was used as a positive control at10 mg/kg and resulted in a transient parasitemia peak of 2.5% between DPI-10 and DPI-15, followed by complete parasite clearance by DPI-17 and 100% survival (Fig. 3B and K). In contrast, none of the prioritized acridone derivatives evaluated *in vivo* significantly reduced parasitemia or improved survival compared to vehicle controls (Figure 3C-J and K) like what was observed for T111 earlier (Fig. 2A, B).

**Figure 3.**
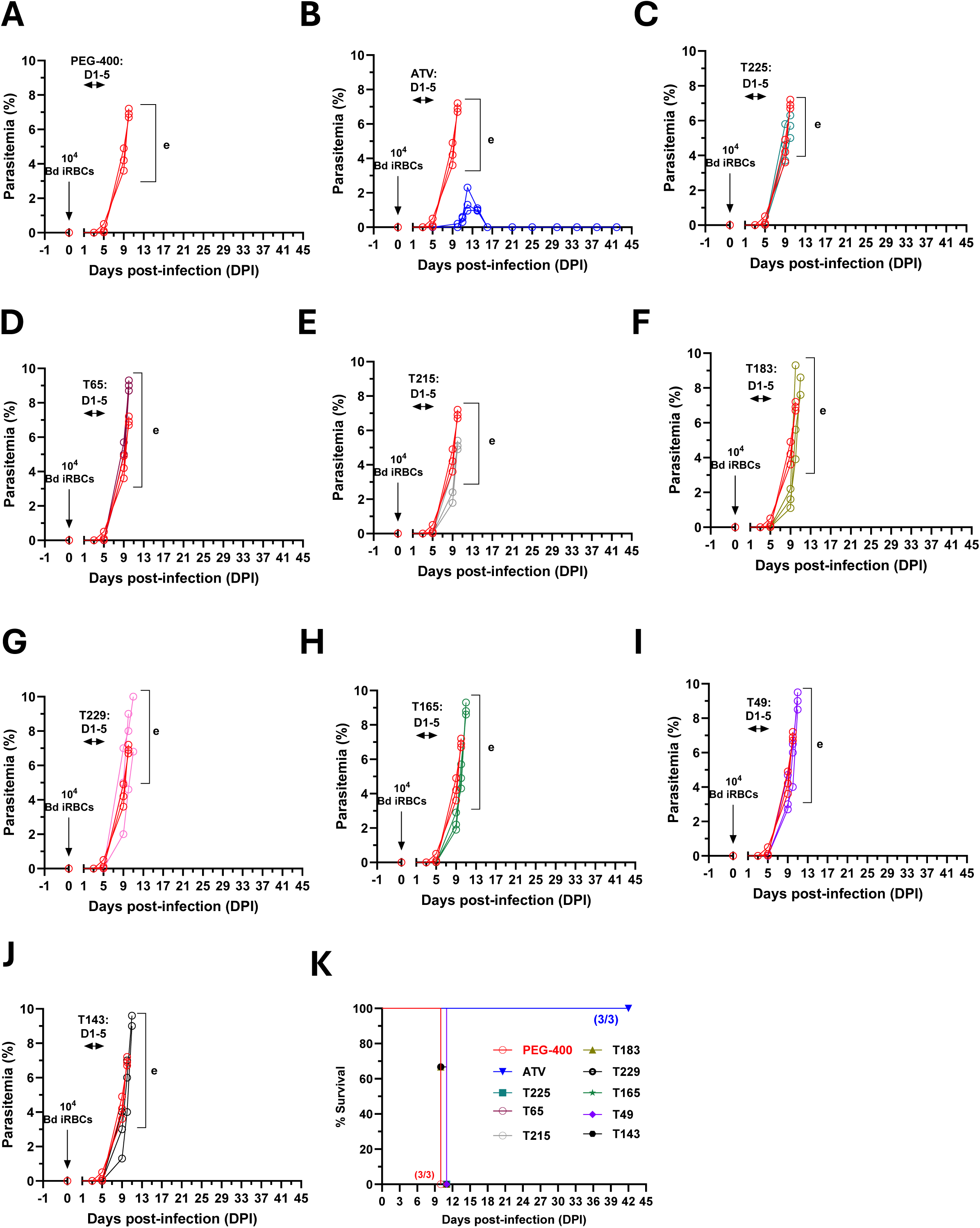
*In vivo* efficacy of eight prioritized acridone compounds against *Babesia duncani* infection in mice. **(A-B)** Parasitemia in female C3H/HeJ mice (n = 3 per group) intravenously infected with 1 × 10^4^ *Babesia duncani* (WA-1 strain)-infected red blood cells (iRBCs), inducing lethal infection. Mice were treated via oral gavage once daily from days post-infection (DPI) 1 to 5 with either (**A**) PEG-400 (vehicle, red) or (**B**) Atovaquone (10 mg/kg blue). **(C-J).** Parasitemia in female C3H/HeJ mice (n = 3 per group) intravenously infected with 1 × 10⁵ *Babesia duncani*-infected red blood cells (iRBCs), inducing lethal infection. Mice were treated via oral gavage once daily from DPI 1 to 5 with prioritized acridone derivatives (30 mg/kg). Parasitemia was quantified daily using Giemsa-stained blood smears, counting a minimum of 3,000 erythrocytes per smear. **(K)** Kaplan-Meier survival curve for female C3H/HeJ mice infected with *B. duncani* and treated with vehicle (PEG-400), Atovaquone (ATV) or eight prioritized acridone derivatives, illustrating survival outcomes over the study period. e, euthanized.

## Discussion

In this study, we provide a detailed evaluation of a diverse panel of acridone derivatives for their activity against the primary etiological agents of human babesiosis. Building on prior work that established acridones as potent multistage antimalarial agents, we demonstrate that these compounds also display robust and broad-spectrum in vitro efficacy against *B. duncani* and *Babesia divergens*. Nine derivatives were prioritized and exhibited excellent potency with favorable selectivity indices. These findings demonstrate that the acridone scaffold, originally optimized for *Plasmodium* parasites, also possesses intrinsic antibabesial activity and thus represents a promising chemical platform for cross-apicomplexan drug development.

Our antiparasitic discovery pipeline leverages a recently established In Culture-In Mouse (ICIM) system along with a pan-antiparasitic screening platform based on standardized culture condition of hemoprotozoan parasites [13, 18, 19]. Using this system, we performed small-scale SAR studies on the 19 acridones derivatives to identify novel antibabesial candidates. Our hypothesis was based on the proven efficacy of acridones against *Plasmodium* spp, and our findings support this rationale [16, 17].

All evaluated acridones demonstrated low micromolar to nanomolar in vitro activity, with IC_50_s ranging from 3 µM (T143) to as low as 0.1 nM (T111) and excellent selectivity, suggesting a favorable therapeutic window. In contrast to the limitations associated with current CDC-recommended therapies (e.g., atovaquone + azithromycin or clindamycin + quinine), which often require prolonged regimens and are associated with significant side effects, acridones present a safer and potentially more effective alternative. Among the acridones examined in this study, T111 was the most potent, with IC₅₀ values of 0.1 ± 0.009 nM and 0.5 ± 0.6 nM against *B. duncani* and *B. divergens*, respectively, and selectivity indices exceeding 15,000. Notably, T111 has already shown preclinical promise in malaria models due to its efficacy [16].

Recent multi-omics studies have highlighted shared biochemical and structural features between *Plasmodium* and *Babesia* parasites [13, 20, 21]. These similarities, particularly in essential pathways such as nucleotide biosynthesis and mitochondrial function, reveal common vulnerabilities [8, 13, 22–25]. Prior success in targeting such conserved pathways (e.g., the cytochrome *bc*_1_ complex (Cytb)) supports a rationale for acridones as broad-spectrum antiparasitic agents. In fact, acridones tested in this study exhibited superior in vitro activity compared to many existing antibabesial drugs, including atovaquone [26].

Mechanistic studies suggest that the acridone analogs target mitochondrial Cytb of malaria parasites [15, 27], implying a broader inhibitory mechanism. Resistance selection studies, functional assays and biochemical analyses in *Babesia* parasites with these compounds are warranted to confirm target engagement and further elucidate the mechanism of action.

Despite the strong *in vitro* potency and high selectivity of the acridone derivatives, based on favorable physicochemical properties, including improved solubility and the availability of prodrug forms led us the *in vivo* efficacy studies in murine models of *B. duncani* and *B. microti* infections revealed limited therapeutic efficacy. Oral administration of short-listed compounds at doses up to 40 mg/kg for five consecutive days did not significantly reduce parasitemia in the case of *B. microti* infection or parasitemia and survival in the case of *B. duncani* infection. In contrast, the reference control, atovaquone, showed marked in vivo efficacy under the same conditions. The lack of oral efficacy against *Babesia* parasites in mice is surprising as the acridone analogs demonstrated excellent oral efficacy against malaria and toxoplasmosis infections in rodent models [15–17, 28]. Notably, several of these acridone analogs achieved complete cures in both asexual blood-stage *Plasmodium yoelii* and liver-stage *P. berghei* infection models in mice, fully consistent with their picomolar *in vitro* antiplasmodial potency [15–17]. This discrepancy between *in vitro* activity and *in vivo* performance in *Babesia* needs to be further investigated to gain deeper understanding of ADME behavior across different mouse strains and infection models to inform rational design of next-generation acridone analogs. Medicinal chemistry optimization aimed at improving solubility, permeability, and metabolic stability will be instrumental in translating *in vitro* potency into meaningful *in vivo* efficacy. In parallel, formulation strategies, including nanoparticle encapsulation or prodrug development, may help overcome current PK limitations. Finally, evaluating these compounds at higher doses (up to 100 mg/kg) or under extended treatment durations (up to 10 days) as well as in combination therapy regimens represents a logical next step to maximize therapeutic potential.

Beyond their immediate therapeutic implications, these findings support a broader principle: antimalarial scaffolds, such as acridones, offer a valuable foundation for antibabesial drug discovery. This study positions acridones as a tractable and versatile platform for future lead optimization and cross-apicomplexan therapeutic development.

## Materials and Methodology

Unless otherwise stated, all chemicals/reagents and solvents were purchased from commercial supplies and used without further purification. The selected target 19 acridone analogs (Table 1) were synthesized in good to excellent yields using streamlined and high-yielding procedures previously developed by the Kelly team [15–17] for rapidly accessing diverse acridone scaffolds. All intermediate and final target compounds were rigorously characterized by NMR and HRMS analyses, ensuring complete structural conformation and purity suitable for biological evaluation. NMR spectra were recorded on a Bruker AMX-400 spectrometer operating at 400 MHz, using CDCl_3_ and DMSO-*d_6_* as solvents at 25 ℃. HRMS data were obtained by electrospray ionization (ESI) on a Vanquish UHPLC/HPLC system coupled to a Q Exactive Orbitrap mass spectrometer operating at a resolution of 35,000. Analytical HPLC was performed on an Agilent 1260 Infinity II LC System using a C8 column (2.1 × 50 mm) using a linear gradient of water/methanol (containing 10 mM ammonium acetate) from 50:50 to 0:100 over 10 min at a flow rate of 0.5 mL/min. The purity of all target compounds was confirmed to be greater than 95%.

### Animal studies ethics statement

Immunocompetent C3H/HeJ female mice were procured from The Jackson Laboratory. All animal experiments strictly adhered to Yale University’s institutional guidelines for the care and use of laboratory animals and were conducted under the approval of a protocol sanctioned by the Institutional Animal Care and Use Committees (IACUC) at Yale University.

### Cytotoxicity Studies

To assess acridone compounds’ cytotoxicity, we performed MTT assays on HepG2, HeLa, HCT-116, and HEK-293 cell lines, adapting Vydyam et al.[13] with minor optimizations. Cells were trypsinized, seeded into 96-well plates (5,000–10,000 cells/well), and incubated for 24 hours at 37°C in 5% CO₂. Medium was replaced with acridone compounds (0.1–100 µM in <0.5% DMSO), tested in triplicates. After 48 hours, cells were washed with PBS, and 100 µL MTT solution (0.5 mg/mL) was added. Following 3-hour incubation, formazan crystals were dissolved in 100 µL DMSO solvent. Absorbance was read at 570 nm using a BioTek Synergy™ H1. Cell viability was calculated relative to controls after background subtraction. Dose-response curves and IC_50_ values were generated using GraphPad Prism (version 9.4). Therapeutic index was calculated as the ratio of cytotoxicity IC_50_ to drug efficacy IC_50_ (from parasite assays). Experiments included three biological replicates for statistical reliability.

### *In Vitro* Drug Efficacy Studies

The *in vitro* efficacy of acridone compounds against Babesia parasites (*B. duncani strain WA-1 and B. divergens strain Rouen87*) was assessed using a fluorescence-based SYBR Green-I assay, following the protocol reported previously [13, 14]. Human red blood cells (hRBCs) infected with *B. duncani or B. divergens* were cultured in DFS20 complete medium [DMEM-F12 (GIBCO, Cat. 11330-032) supplemented with 20% heat-inactivated fetal bovine serum (GIBCO, Cat. No. A52568-01), 1X HT (Sigma, Cat. no. H0137), 1X Antibiotic/Antimycotic (GIBCO, Cat. No. 15240062) and 1% of 10 mg/ml gentamicin (GIBCO, Cat. No. 15710-064)] at 0.2% parasitemia and 5% hematocrit. Test compounds were either applied at a fixed concentration (1 µM) for initial screening or as a series of 2-fold dilutions (from 10 µM to 0.00119 nM for *B. duncani*, and 100 μM to 0.0119 nM for *B. divergens*) for dose-response studies. Compounds were diluted in DFS20, and 100 µL of the drug-parasite mixture was seeded into 96-well tissue culture plates. Plates were incubated for 62 hours at 37°C in a low-oxygen environment (5% CO₂, 2% O₂, 93% N₂) to mimic physiological conditions. Post-incubation, parasite growth was quantified using the SYBR Green-I assay. An equal volume (100 µL) of SYBR Green-I lysis buffer (0.008% saponin, 0.08% Triton X-100, 20 mM Tris-HCl (pH 7.5), 5 mM EDTA, and 1× SYBR Green-I) was added to each well. Plates were incubated at 37°C for 15 minutes in the dark to facilitate parasite lysis and DNA intercalation. Fluorescence was measured using a BioTek Synergy™ Mx microplate reader with excitation at 480 nm and emission at 580 nm. Background fluorescence from uninfected hRBCs in DFS20 was subtracted to normalize the data. Dose-response curves were plotted using GraphPad Prism (version 9.4), and IC_50_ values were calculated from sigmoidal fits of fluorescence intensity versus drug concentration. Each experiment included pairs of technical replicates and was repeated across three biological replicates to yield mean IC_50_ values ± standard deviation (SD). Positive controls (atovaquone at 1 µM) and negative controls (vehicle-only) were included in each plate to validate assay performance.

### *In Vivo* Drug Efficacy Studies in Murine Models

The *in vivo* efficacy of acridone compounds against *B. duncani* (strain WA-1) and *B. microti* (strain LabS1) was evaluated in C3H/HeJ mice following the protocol reported previously [14, 18, 19]. Female mice (5–6 weeks old) were housed under specific pathogen-free conditions. Groups of 3–5 mice were infected intravenously with an inoculum of either 10⁴ or 10⁵ infected RBCs (*B. duncani or B. microti* (as indicated) to establish infection. Treatment was initiated either on day 1 post-infection (DPI 1) or DPI 3 and continued daily for 5 days (until DPI 5 or DPI 7, respectively). Acridone compounds were administered via oral gavage at doses of 30 mg/kg or 40 mg/kg, formulated in 100 µL of PEG-400 as the vehicle. Control groups received either vehicle alone (PEG-400) or atovaquone (10 mg/kg) as a positive control. The treatment volume was standardized at 100 µL per mouse to minimize stress and ensure consistent delivery. Parasitemia was monitored by collecting tail vein blood samples (5–10 µL) at predetermined time points (e.g., DPI 3, 5, 7, and 10). Blood smears were prepared, fixed with methanol, and stained with Giemsa. Parasitemia was quantified by light microscopy, counting a minimum of 3,000 erythrocytes per smear to ensure accuracy. Data were expressed as the percentage of infected RBCs. Body weight and clinical signs (e.g., lethargy, ruffled fur) were monitored daily to assess drug tolerability.

## Acknowledgement, Funding

We gratefully acknowledge the financial support provided by the National Institutes of Health (NIH) through grants AI123321, AI138139, AI152220, and AI136118, to C.B.M research laboratory, which enabled this research. We also thank our colleagues and collaborators for their valuable insights and technical assistance throughout the study. Synthesis of acridone compounds was supported by the NIH/NIAID (award AI158533), DOD/PRMRP (award W81XWH2210494), and VA BLRD (award 1I01BX005674).

## Data Availability

All data generated or analyzed during this study are included in this published article and its supplementary information files.

## Ethics Approval

Human blood (A^+^ Packed RBCs), donated by anonymized individuals, was obtained from the American Red Cross under approval account number N0337833H.

## References

1. Krause, P.J., et al., Clinical Practice Guidelines by the Infectious Diseases Society of America (IDSA): 2020 Guideline on Diagnosis and Management of Babesiosis. Clin Infect Dis, 2021. 72(2): p. e49–e64.

2. Swanson M, P.A., Williamson J, Montgomery S., Trends in Reported Babesiosis Cases — United States, 2011–2019. 2023.

3. Krause, P.J., Human babesiosis. Int J Parasitol, 2019. 49(2): p. 165–174.

4. Herwaldt, B., et al., A fatal case of babesiosis in Missouri: identification of another piroplasm that infects humans. Ann Intern Med, 1996. 124(7): p. 643–50.

5. Scott, J.D., et al., Detection of Babesia odocoilei in Humans with Babesiosis Symptoms. Diagnostics (Basel), 2021. 11(6).

6. Herwaldt, B.L., et al., Babesia divergens-like infection, Washington State. Emerg Infect Dis, 2004. 10(4): p. 622–9.

7. Vannier, E., B.E. Gewurz, and P.J. Krause, Human babesiosis. Infect Dis Clin North Am, 2008. 22(3): p. 469–88, viii-ix.

8. Renard, I. and C. Ben Mamoun, Treatment of Human Babesiosis: Then and Now. Pathogens, 2021. 10(9).

9. Krause, P.J., et al., Persistent and relapsing babesiosis in immunocompromised patients. Clin Infect Dis, 2008. 46(3): p. 370–6.

10. Vannier, E.G., et al., Babesiosis. Infect Dis Clin North Am, 2015. 29(2): p. 357–70.

11. Wormser, G.P., et al., Emergence of resistance to azithromycin-atovaquone in immunocompromised patients with Babesia microti infection. Clin Infect Dis, 2010. 50(3): p. 381–6.

12. Simon, M.S., et al., Clinical and Molecular Evidence of Atovaquone and Azithromycin Resistance in Relapsed Babesia microti Infection Associated With Rituximab and Chronic Lymphocytic Leukemia. Clin Infect Dis, 2017. 65(7): p. 1222–1225.

13. Vydyam, P., et al., In vitro efficacy of next-generation dihydrotriazines and biguanides against babesiosis and malaria parasites. Antimicrob Agents Chemother, 2024. 68(9): p. e0042324.

14. Vydyam, P., et al., Tafenoquine-Atovaquone Combination Achieves Radical Cure and Confers Sterile Immunity in Experimental Models of Human Babesiosis. J Infect Dis, 2024. 229(1): p. 161–172.

15. Dodean, R.A., et al., Discovery and Structural Optimization of Acridones as Broad-Spectrum Antimalarials. J Med Chem, 2019. 62(7): p. 3475–3502.

16. Dodean, R.A., et al., Development of Next-Generation Antimalarial Acridones with Radical Cure Potential. J Med Chem, 2025. 68(8): p. 8817–8840.

17. Kancharla, P., et al., Lead Optimization of Second-Generation Acridones as Broad-Spectrum Antimalarials. J Med Chem, 2020. 63(11): p. 6179–6202.

18. Pal, A.C., et al., Babesia duncani as a model organism to study the development, virulence and drug susceptibility of intraerythrocytic parasites in vitro and in vivo. J Infect Dis, 2022.

19. Kumari, V., et al., Babesia duncani in Culture and in Mouse (ICIM) Model for the Advancement of Babesia Biology, Pathogenesis, and Therapy. Bio Protoc, 2022. 12(22).

20. Singh, P., et al., Babesia duncani multi-omics identifies virulence factors and drug targets. Nat Microbiol, 2023. 8(5): p. 845–859.

21. Singh, P., et al., Insights into the evolution, virulence and speciation of Babesia MO1 and Babesia divergens through multiomics analyses. Emerg Microbes Infect, 2024. 13(1): p. 2386136.

22. Lawres, L.A., et al., Radical cure of experimental babesiosis in immunodeficient mice using a combination of an endochin-like quinolone and atovaquone. J Exp Med, 2016. 213(7): p. 1307–18.

23. Chiu, J.E., et al., Effective Therapy Targeting Cytochrome bc(1) Prevents Babesia Erythrocytic Development and Protects from Lethal Infection. Antimicrob Agents Chemother, 2021: p. Aac0066221.

24. Chiu, J.E., et al., Cytochrome b Drug Resistance Mutation Decreases Babesia Fitness in the Tick Stages But Not the Mammalian Erythrocytic Cycle. The Journal of Infectious Diseases, 2021. 225(1): p. 135–145.

25. Dews, E.A., et al., A propidium iodide-based in vitro screen of the “Bug Box” against Babesia duncani reveals potent inhibitors. Antimicrob Agents Chemother, 2025. 69(7): p. e0003525.

26. Abraham, A., et al., Establishment of a continuous in vitro culture of Babesia duncani in human erythrocytes reveals unusually high tolerance to recommended therapies. J Biol Chem, 2018. 293(52): p. 19974–19981.

27. Kancharla, P., et al., Potent Acridone Antimalarial against All Three Life Stages of Plasmodium. Res Sq, 2025.

28. Alday, P.H., et al., Acridones Are Highly Potent Inhibitors of Toxoplasma gondii Tachyzoites. ACS Infect Dis, 2021. 7(7): p. 1877–1884.

